# Modeling Single Cell Trajectory Using Forward-Backward Stochastic Differential Equations

**DOI:** 10.1101/2023.08.10.552373

**Authors:** Kevin Zhang, Junhao Zhu, Dehan Kong, Zhaolei Zhang

## Abstract

Recent advances in single-cell sequencing technology have provided opportunities for mathematical modeling of dynamic developmental processes at the single-cell level, such as inferring developmental trajectories. Optimal transport has emerged as a promising theoretical framework for this task by computing pairings between cells from different time points. However, optimal transport methods have limitations in capturing nonlinear trajectories, as they are static and can only infer linear paths between endpoints. In contrast, stochastic differential equations (SDEs) offer a dynamic and flexible approach that can model non-linear trajectories, including the shape of the path. Nevertheless, existing SDE methods often rely on numerical approximations that can lead to inaccurate inferences, deviating from true trajectories. To address this challenge, we propose a novel approach combining forward-backward stochastic differential equations (FBSDE) with a refined approximation procedure. Our FBSDE model integrates the forward and backward movements of two SDEs in time, aiming to capture the underlying dynamics of single-cell developmental trajectories. Through comprehensive benchmarking on multiple scRNA-seq datasets, we demonstrate the superior performance of FBSDE compared to other methods, highlighting its efficacy in accurately inferring developmental trajectories.

## 1 Introduction

The technology of single-cell RNA sequencing (scRNA-seq) is a revolutionary breakthrough in the study of cellular developmental dynamics [1, 2]. With the availability of time series scRNA-seq data, lineage tracing and cell differentiation processes have been studied in multiple cell types such as salivary glands, liver cells, lung cells, kidney cells, neuronal cells, and tumor cells [3–8]. Despite their success, the sequencing process is destructive as the sampled cells are destroyed and their future gene expressions cannot be measured. Several mathematical approaches have emerged to take these time series data as input and recover the true cellular differentiation processes. Earlier modelling approaches include continuous-state hidden Markov models (CSHMM) [9], and a graph-based model Tempora [10]. These methods typically cluster cells into a small number of distinct cell types and model the trajectory of these cell types instead of modelling individual cells. These approaches are simpler to model and are easier to interpret but lose cell-level information after reducing a population of cells into discrete cell types. Recently, optimal transport (OT) becomes a state-of-art approach in modelling cell developmental trajectories from single cell RNA-seq data. In particular, Schiebinger [11] described a new framework (Waddington-OT) to identify the pairings between cells from different time points where gene expressions of single cells are assumed to follow a developmental trajectory with the lowest transport energy cost. Waddington-OT uses a joint probability mass function to infer how likely a cell sampled at a previous time point will become another cell sampled at a later time point. The key of the OT model is that it assumes the cell population evolves in an energy-efficient fashion analogous to the optimal transport plan in probability theory. In this context, the energy term is defined as the differences in genome-wide gene expression profile of a population of cells between consecutive time points [11]. In contrast to graph-based models, optimal transport does not require clustering cells into discrete cell types. Instead, it models the developmental processes at the single cell level, making it a potentially more powerful approach.

Despite its success and later improvement [12], Waddington-OT still has two limitations. First, it can only infer the endpoints of a path from sampled cells, making generative modelling infeasible because paths starting from an unobserved point cannot be simulated. Second, path or trajectory inference in Waddington-OT is static as it only provides linear interpolation between endpoints. In this context, linear trajectory means the shape of a path is simply a straight line connecting the endpoints. Instead of matching path endpoints with OT, a potential solution is to model path movements with stochastic differential equations (SDEs), which are more adept in dynamic trajectory inferences [13, 14]. In general, an SDE model considers *X*_*t*_ as the gene expression of a cell in a population existing between initial time *t* = 0 and terminal time *t* = *T* ; the change in *X*_*t*_ at time *t* is denoted by *dX*_*t*_ and equals *vdt*+*σdW*_*t*_ where *v* and *σ* represent drift and volatility. The drift term *v* takes a cell’s spatial and temporal information as input and produces the direction in which the cell changes its gene expression as output. The volatility term *σ* mimics the pure randomness in cell movements by scaling Brownian motions denoted by *W*_*t*_. As a result, SDE is able to parametrize non-linear trajectories and provide feasible generative simulations on a continuous timeline. The volatility term *σ* in SDE is also equivalent to the entropy regulation in optimal transport [15]. In practice, SDE models can be parameterized by neural networks [16, 17], which have shown good performance in recovering scRNA trajectories [18].

Single cell sequencing has allowed us to study cell-to-cell interactions, which has not been included in earlier trajectory inference methods [19, 20]. In biology, it is well-established that cells can influence the development of other cells by modulating their gene expression or other cellular programs. This modulation can occur through the secretion of small molecules or via direct interactions between receptors and ligand molecules on cell surfaces. Interestingly, these cell-to-cell interactions bear resemblance to concepts found in mean field control theory, which aims to optimize the collective behavior of a population while accounting for interactions among individual members [16, 17]. Drawing inspiration from this, a recent model called TrajectoryNet [18] incorporated mean field control theory to capture interactions among single cells.

In the following, we explore the rationale and limitations of TrajectoryNet, which is the most closely related method to our proposed approach. It serves as a valuable baseline for our method. The model assumes that gene expressions in cells at each time point *t* = 0, 1, 2, …, *T* follow a distribution *q*_*t*_. TrajectoryNet begins with *q*_0_ and generates a sequence of distributions *ρ*_*t*_ for *t* = 1, 2, …, *T* based on the SDE model (refer to **Equation** 1 in Section 4). This process can be achieved through simulations, starting with *m* cells randomly subsampled from time point *t* = 0 and then simulating subsequent cells at time points *t* = 1, 2, …, *T* using the SDE model. The gene expressions of these simulated cells are denoted by 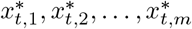, and their empirical distribution is denoted as 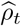. The main optimization problem in TrajectoryNet is similar to a road-building problem, where the objective is to construct a road connecting two points *A* and *B*, while passing through multiple intermediate checkpoints. TrajectoryNet aims to find an optimal SDE model that captures the collective displacement of simulated cells, akin to the construction cost of a road. Additionally, TrajectoryNet incorporates interactions among cells, resembling the movement of cars passing and merging on the road. Similar to the road-building problem, one would expect 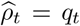 for each time point *t* = 0, 1, 2, …, *T*, which can be interpreted as the initial and terminal points of the road and the intermediate checkpoints it must traverse. However, TrajectoryNet only satisfies the initial constraint 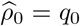 but not the subsequent constraints at *t* = 1, 2, …, *T* . Instead, TrajectoryNet relaxes these constraints by penalizing the divergence between *q*_*t*_ and 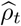. In other words, TrajectoryNet approximates the constraints as 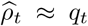 for *t* = 1, 2, …, *T* . Consequently, TrajectoryNet is less effective due to the lack of exact matching between the simulated distributions 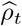 and the true distributions *q*_*t*_ for *t* = 1, 2, …, *T* .

Motivated to overcome these limitations, we propose two key enhancements to improve trajectory inference. Firstly, we enhance the approximation of intermediate constraints by introducing a new penalty term in the optimization problem. Secondly, we enforce the terminal constraint 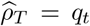 by implementing the FBSDE method (refer to **Figure** 1 for a visual illustration). The proposed enhancements aim to enhance the accuracy of trajectory inference by achieving a closer alignment between the simulated distributions 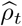 and the true distributions *q*_*t*_ for each time point *t* = 1, 2, …, *T* . This improved alignment ensures that the inferred trajectories more faithfully capture the underlying dynamics of the system. In the rest of this section, we provide an overview of our innovative approaches and their key components.

**Fig. 1:**
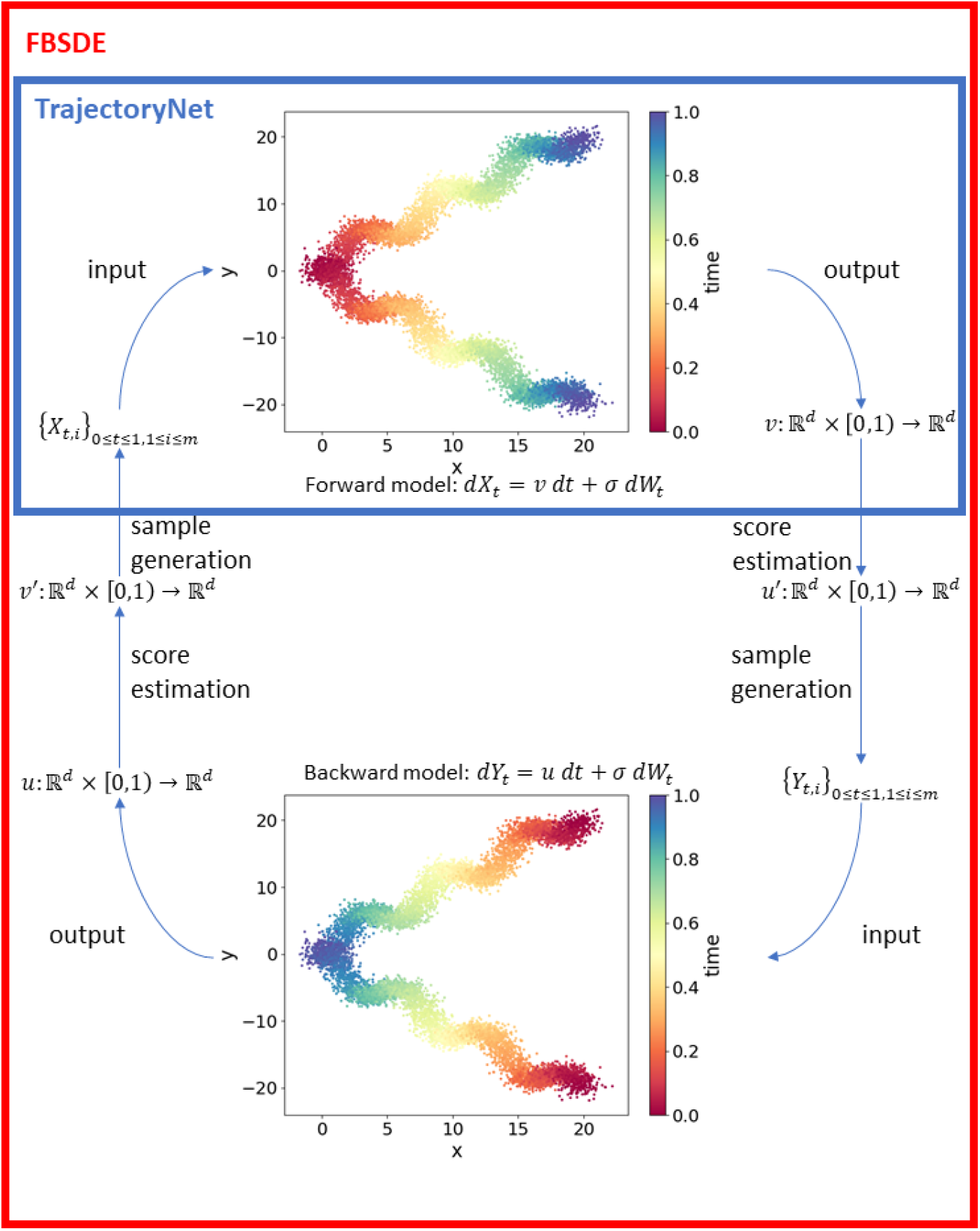
Overview of FBSDE model. Colours in the scatter plots represent time where red is the earliest and blue is the latest time point. In the forward model, the population is modelled from left to right. In the backward model, the population evolves from right to left. The blue rectangle on top represents the TrajectoryNet framework by Tong et al [18] which is equivalent to half of one iteration of our FBSDE model. The FBSDE model iterates between the Forward and Backward models; traversing through the Forward model generates new simulated data points which are subsequently used as training set by the backward model and vice versa.

To enhance the approximation of intermediate constraints, we introduce a new penalty term as compared to the TrajectoryNet method discussed above [18]. To satisfy the conditions 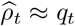 for all time points *t*, TrajectoryNet implements two penalty terms. First, it directly penalizes the divergence between *p*_*t*_ and 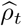 using the maximum likelihood function. Second, it also introduces a penalty term which is proportional to the sum of distances among *k* nearest neighbours. To illustrate this, suppose we have *n* observations and *n* simulations at each time point *t* and denote *x*_*t,i*_ and 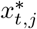 the gene expressions of *i*th observed cell and *j*th simulated cell respectively at time *t*. The conditions 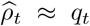 imply that there is a large overlap between the locations occupied by the observations and those by the simulations in the gene expression space (see **Figure** 2a-2b). At each time point *t*, for each simulated data point 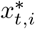 (i.e., gene expression of single cells), TrajectoryNet computes its distances to *k* closest observations among {*x*_*t,i*_}. These soft marginal constraints force the simulated points to be in a close neighbourhood of observations. In other words, it is less likely to have simulations far away from observations. Therefore, this penalty improves the precision of the model. We modified the k-nearest-neighbour distances in TrajectoryNet [18] by additionally computing the distances to *k* closest simulations among 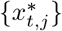 for each observed point *x*_*t,i*_. This additional term allows more observations to be included in a close neighbourhood of simulated points (see Section 4 for details). As a result, our model also improves recall as illustrated in **Figure** 2b.

**Fig. 2:**
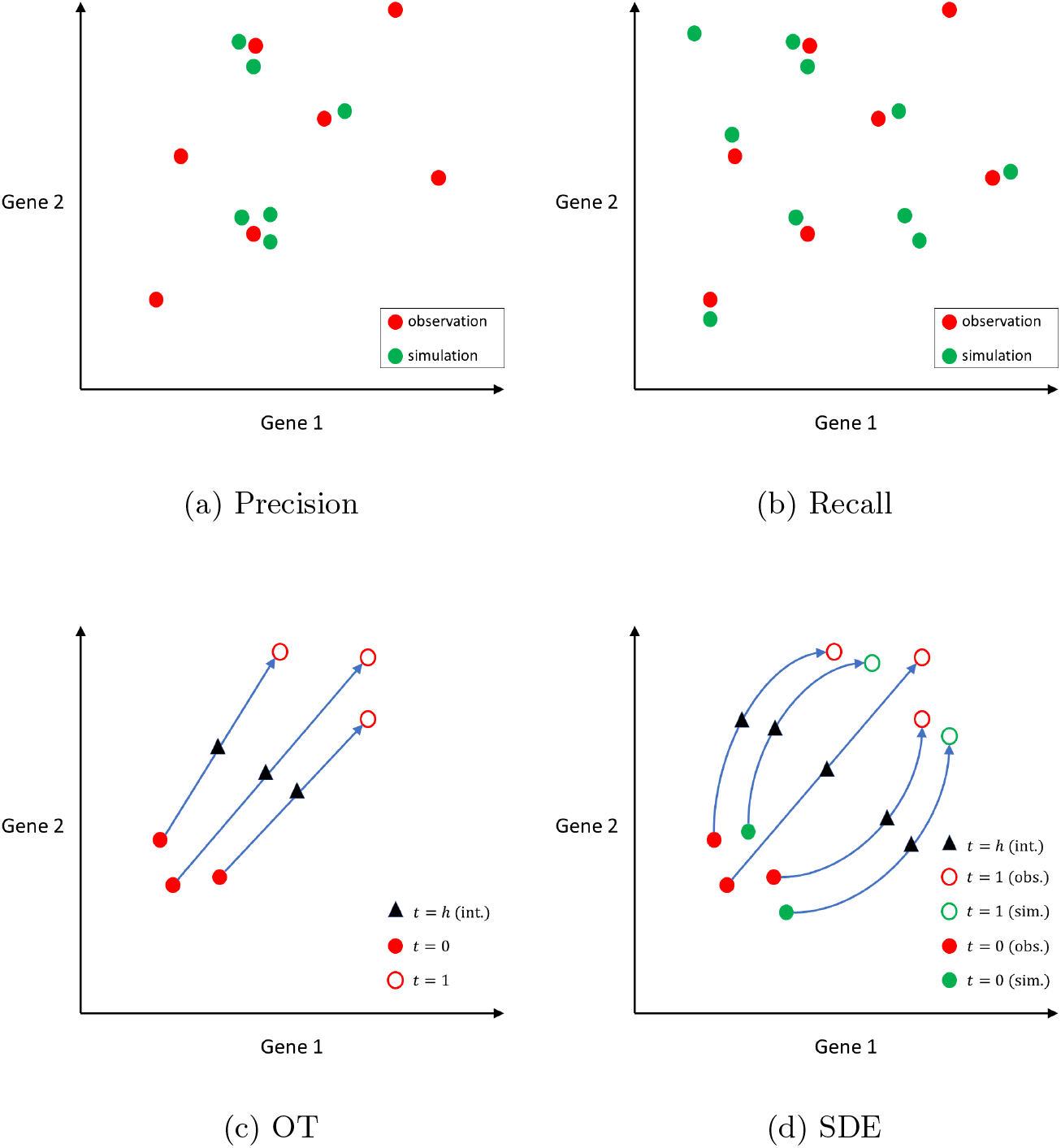
2a-2b) Comparison between precision and recall. Red circles represent observations and green circles represent simulations. In 2a, a high precision model means that simulations are close to observations; however, there can be some observations without any simulations nearby. In 2b, a high recall model means that most observations have some simulations nearby; however, there can be some simulations that are far away from any observations. 2c-2d) Comparison between OT and SDE. Filled circles represent cells from *t* = 0. Unfilled circles represent cells from *t* = 1. The red colour represents observations. The green colour represents simulations. Triangles indicated interpolated cells at time *t* = *h*. 2c) OT infers the endpoints of a path and the trajectory is simply a straight line connecting the endpoints. OT can only infer paths whose endpoints come from observations. 2d) SDE infers both linear and non-linear trajectories using differential equations. In addition, SDE also infers paths originating from points that are not observations (i.e., given an arbitrary starting point).

To enforce the exact terminal equality constraint 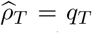, we utilize the forward-backward stochastic differential equations (FBSDE) proposed by Vargas et al [21]. The FBSDE model consists of a forward model which goes from initial time to terminal time (0 → *T*) and a backward model which goes from terminal time to initial time (*T →* 0) (see Section 4 for details). The forward model generates training data for the backward model and vice versa. The core concept of FBSDE is that training the two models where the time goes in opposite directions will eventually lead to convergence such that 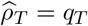 is met by the forward model and 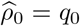 is met by the backward model. A recent study by Bunne et al models cell trajectory with FBSDEs and demonstrates that the model indeed preserves terminal equality constraints 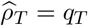 [22]. However, this implementation of the FBSDE model considers only the initial and terminal constraints, and does not model the intermediate time points. To further utilize the information from these observed intermediate time points, we extend the original FBSDE by enforcing soft constraints along all the intermediate time points as discussed above.

We benchmark FBSDE with two other methods Waddington-OT (WOT) and TrajectoryNet on three scRNA data sets: Human Embryonic Stem Cells Dataset [23], Mouse Embryonic Fibroblasts Dataset [11] and *Arabidopsis thaliana* Stem Cells Dataset [12]. We adopt a standard leave-one-out benchmark framework (i.e., removing one time point and trying to recover gene expression of hidden time point from other time points). Our results show that the FBSDE model consistently outperforms TrajectoryNet and Waddington-OT.

## 2 Results

### 2.1 Overview of the FBSDE model

**Figure** 1 provides an overview of the FBSDE. The key innovation of our approach over previously published methods such as TrajectoryNet [18] is that we model the population growth in both forward and backward directions. As such, the FBSDE framework consists of two parts, the forward model and the backward model; the forward model is essentially the same as TrajectoryNet. It has been widely observed that using Forward model alone may not always satisfy the terminal equality constraints if the true trajectory has many non-linear segments [21]. This can be attributed to the lack of closed form solutions in modelling stochastic differential equations [22]. In other words, the stochastic differential equations cannot be solved analytically. As an alternative, numerical approximations sometimes can alleviate these difficulties, but their performances are not always satisfactory. By including a Backward model in the FBSDE framework, one can convert the terminal constraint 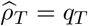 to an initial constraint for the Backward model which can be numerically satisfied [21]. Furthermore, the iteration between the Forward model and the Backward model has been proven to converge to an optimal solution where both the initial constraint 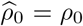 and the terminal constrain 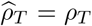 are both satisfied [24].

As alluded to in the Introduction, optimal transport (OT) has been recently adopted in modelling cell growth trajectory and achieved promising performance [11, 12]. Typically, OT is more effective in modelling linear trajectories, whereas SDE can model non-linear trajectories. **Figure** 2c-2d illustrate the differences between OT and SDE. An OT model connects the endpoints with a straight line. In contrast, an SDE model infers non-linear trajectories connecting the endpoints. In addition, an OT model (**Figure** 2c) can only infer trajectories whose endpoints are observations whereas an SDE model (**Figure** 2d) can generate trajectories originating from any arbitrary starting point. Figures 2c through 2d display the trajectories of the cells from time *t* = 0 to *t* = 1 and demonstrate how to interpolate their gene expression at some intermediate time point *t* = *h*, where 0 *< h <* 1. The generative modelling differences between OT and SDE are further illustrated in **Figure** 3. When given a set of initial and terminal distributions, OT can only predict a linear trajectory, as depicted in Figure 3E. On the other hand, SDE can infer trajectories with various shapes depending on the parameter choices, as shown in **Figure** 3a-3d. Specifically, the shapes of the trajectories are affected by the extent to which the population accommodates high entropy (i.e. more crowded), as indicated in this example. As shown in **Figure** 3a-3d, the paths display greater angular rotation as the model is trained to accommodate higher levels of entropy. This is reflected in the curly shape of the paths, which allows the population to remain denser for longer periods. In contrast, optimal transport employs linear path interpolations as shown in **Figure** 3e.

**Fig. 3:**
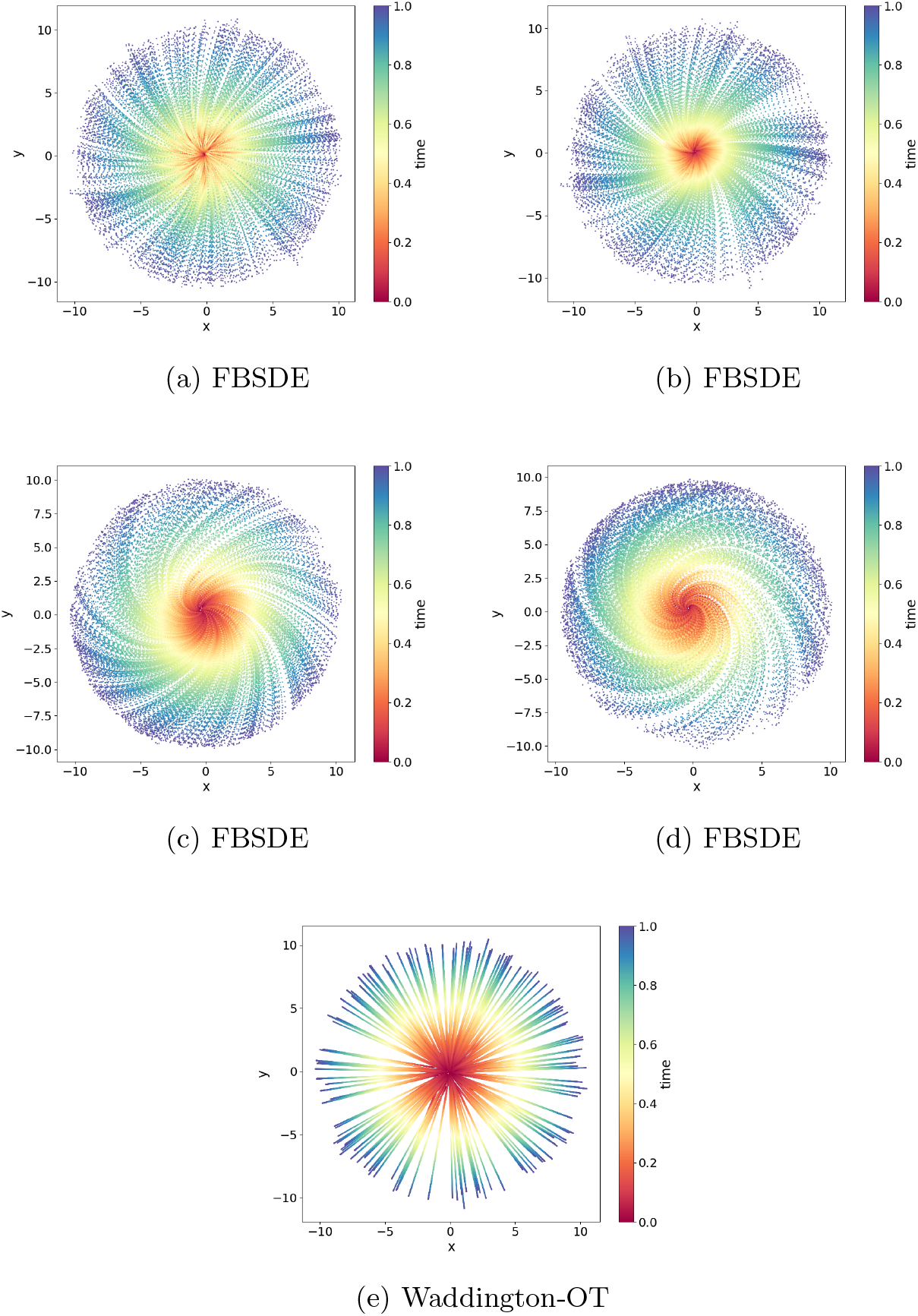
Difference between FBSDE and Waddington-OT in modelling a simulated dataset. The time is coded in the vertical bars next to each panel; the time points are normalized with 0 and 1 representing starting and end point, respectively. At *t* = 0, the population follows the standard normal distribution. At *t* = 1, the population is uniformly distributed on the ring centred at (0, 0) with a radius of 10. At 0 *< t <* 1, the distribution is interpolated by either the FBSDE or Waddington-OT model. In 3a, 3b, 3c and 3d, the population evolves based on a stochastic differential equation where a neural network parametrizes the drift term. From 3a to 3d, the population exhibits more angular rotation as it favours a higher level of entropy (i.e. density). In 3e, the endpoints for each point at *t* = 0 are sampled based on the optimal transport map and the interpolation is performed linearly by connecting each pair of points.

### 2.2 Performance on a Human Embryonic Stem Cells Dataset

We compared the FBSDE model with WOT and TrajectoryNet by visually examining the generative power using two-dimensional human embryonic stem cells data obtained from Moon et al [23]. The dataset contained 16, 825 cells grown as embryoid bodies, which were sampled from 10 time points over a period of 27 days at 3-day intervals. All three models were provided with the same starting population of cells at the initial time point (*t* = 0) and generated samples at 200 equally spaced time points in addition to the already observed time points. **Figure** 4 illustrates the comparison among the ground truth distribution and samples generated by all three models, demonstrating that our FBSDE model generated samples that more closely resemble the observations from human embryonic stem cells data.

**Fig. 4:**
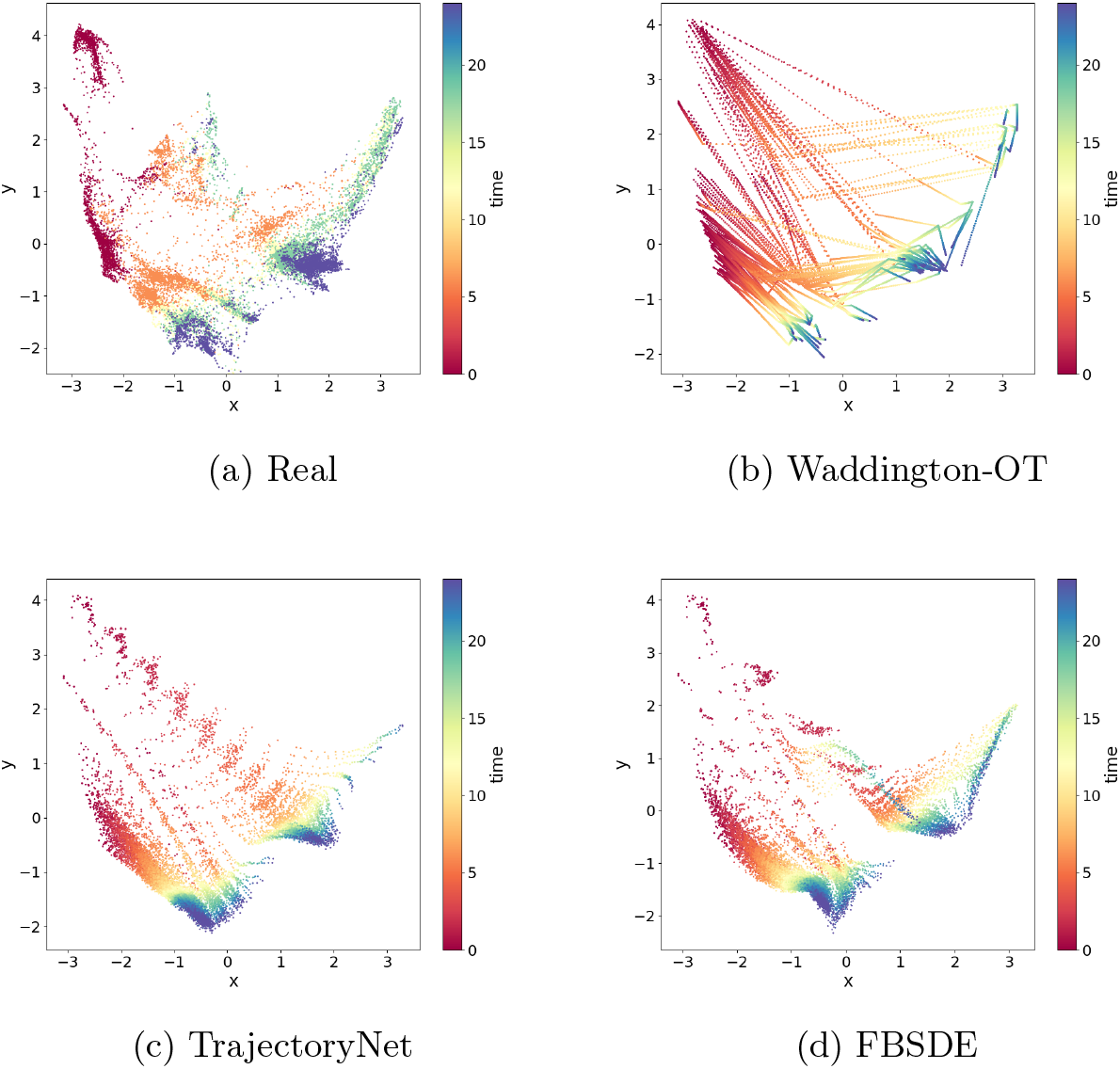
Performances on a human embryonic stem cells dataset. 4a) Gene expression profiles of single cells are reduced to two dimensions using the PHATE method. The data correspond to a total of 10 time points over 27 days at 3 days intervals. The time points are color coded on a spectrum from red to deep blue. 4b-4d) Trajectory inference by three different methods (Waddington-OT, TrajectoryNet and FBSDE). In each experiment, a total of 200 equally spaced time points are added between the starting and end time points. As can be seen, FBSDE provides a trajectory most resembling the ground truth in 4a.

Furthermore, **Figure** 4b highlights why linear interpolations can fail in trajectory inference. The WOT model interpolated some points mainly in the upper portion of the figure that were not seen in real observations. Figures 4c and 4d show that both TrajectoryNet and FBSDE generated simulations that seemed to match the observations at the terminal time point *t* = *T* . In addition, both TrajectoryNet and FBSDE generate smooth trajectories as seen in the observations (**Figure** 4a) in contrast to the segmented linear trajectories generated by the WOT model. However, at intermediate time points, the FBSDE generated simulations represented the observations more accurately than TrajectoryNet. Finally, the FBSDE model visually resembled the observations more closely than the other two models, albeit by a small margin.

### 2.3 Performance on a Mouse Embryonic Fibroblasts Dataset

We further evaluated the generative power of FBSDE, WOT, and TrajectoryNet using a mouse embryonic fibroblasts dataset obtained from Schiebinger et al [11]. This dataset contains 259, 155 cells sampled at 36 time points over an 18-day period. **Figure** 5 shows that all three models performed relatively well on this dataset. Notably, the observations in **Figure** 5a exhibit two distinct sharp turns (indicated by arrows in **Figure** 5) in the shape, which were effectively captured by both WOT (**Figure** 5b) and FBSDE (**Figure** 5d) models but not recognized by TrajectoryNet (**Figure** 5c). However, unlike WOT, which relies on simulations using observed points (as depicted in **Figure** 2c), FBSDE utilizes a neural network to fully parameterize the population movement. FBSDE is a mesh-free method, which means that once a model has been fitted, simulations can be generated without relying on observations. This demonstrates that the FBSDE model has the ability to capture not only smooth but also sharp shape features of developmental trajectories. The successful performance of FBSDE on this dataset provides further evidence of its effectiveness in accurately modelling complex biological processes. Furthermore, a thorough evaluation of the numerical performances of these three methods will be conducted using cross-validation, which will be discussed in detail in Section 2.5.

**Fig. 5:**
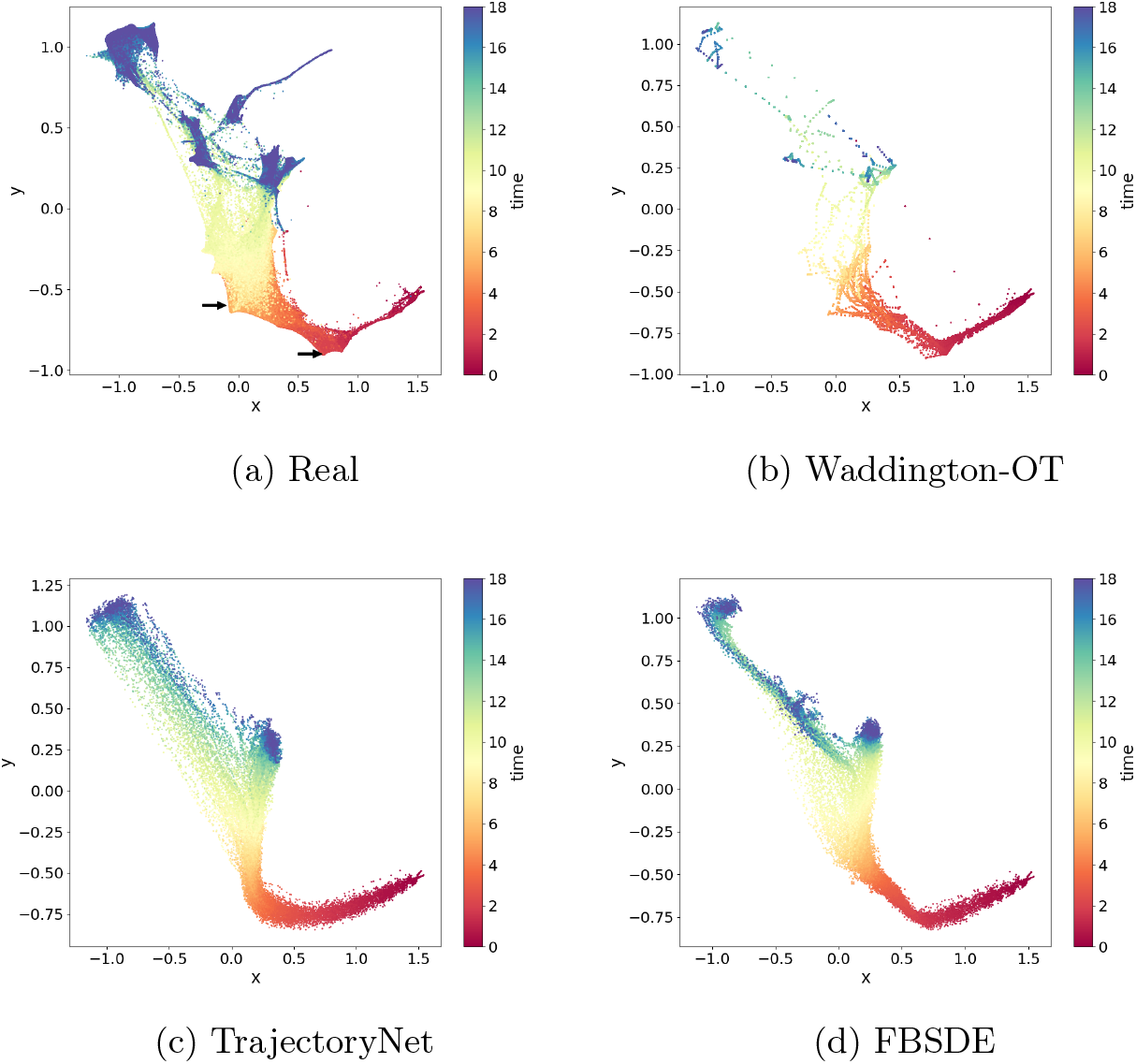
Performance on a mouse embryonic fibroblasts dataset. 5a) Gene expression of single cells are reduced to two dimensions using the force-directed layout embedding proposed in the study by Weinreb and colleagues [25]. The data correspond to a total of 37 time points over 18 days at 12 hours intervals. The time points are color coded on a spectrum from red to deep blue. Two visible sharp turns are indicated by arrows. 5b-5d) Trajectory inference by three different methods (Waddington-OT, TrajectortNet and FBSDE). In each experiment, a total of 200 equally spaced time points are added between the starting and end time points. The same color code is used. As can be seen, all three methods provide trajectories similar to the ground truth in 5a although FBSDE outperforms the others modestly.

### 2.4 Performance on a *Arabidopsis thaliana* Stem Cells Dataset

Finally, we compared the performance of FBSDE, WOT, and TrajectoryNet on the Arabidopsis thaliana stem cells dataset generated by Shahan and colleagues [26]. This dataset contains 110, 427 cells with pseudotime labels ranging from 0 to 50 and exhibits several distinct developmental branches. **Figure** 6a illustrates the complexity of this dataset, making it the most challenging to model compared to the previous two datasets. **Figure** 6c shows that TrajectoryNet simulations exhibit a significant visual discrepancy from the observations in **Figure** 6a. The simulated distributions at observed time points do not match the observed data points, reflecting the challenging nature of the dataset. The WOT model simulations in **Figure** 6b resemble the observations at observed time points, but there appear to be many crossings among branches that are not likely to be biologically relevant. In contrast, the FBSDE simulations in **Figure** 6d resemble the observations more closely, with a few recognizable branches and almost no unrealistic long-range crossings seen. In summary, none of the models perform well on the Arabidopsis thaliana stem cells dataset. However, the WOT model retains shape at observed time points, while the FBSDE model is capable of distinguishing some developmental branches.

**Fig. 6:**
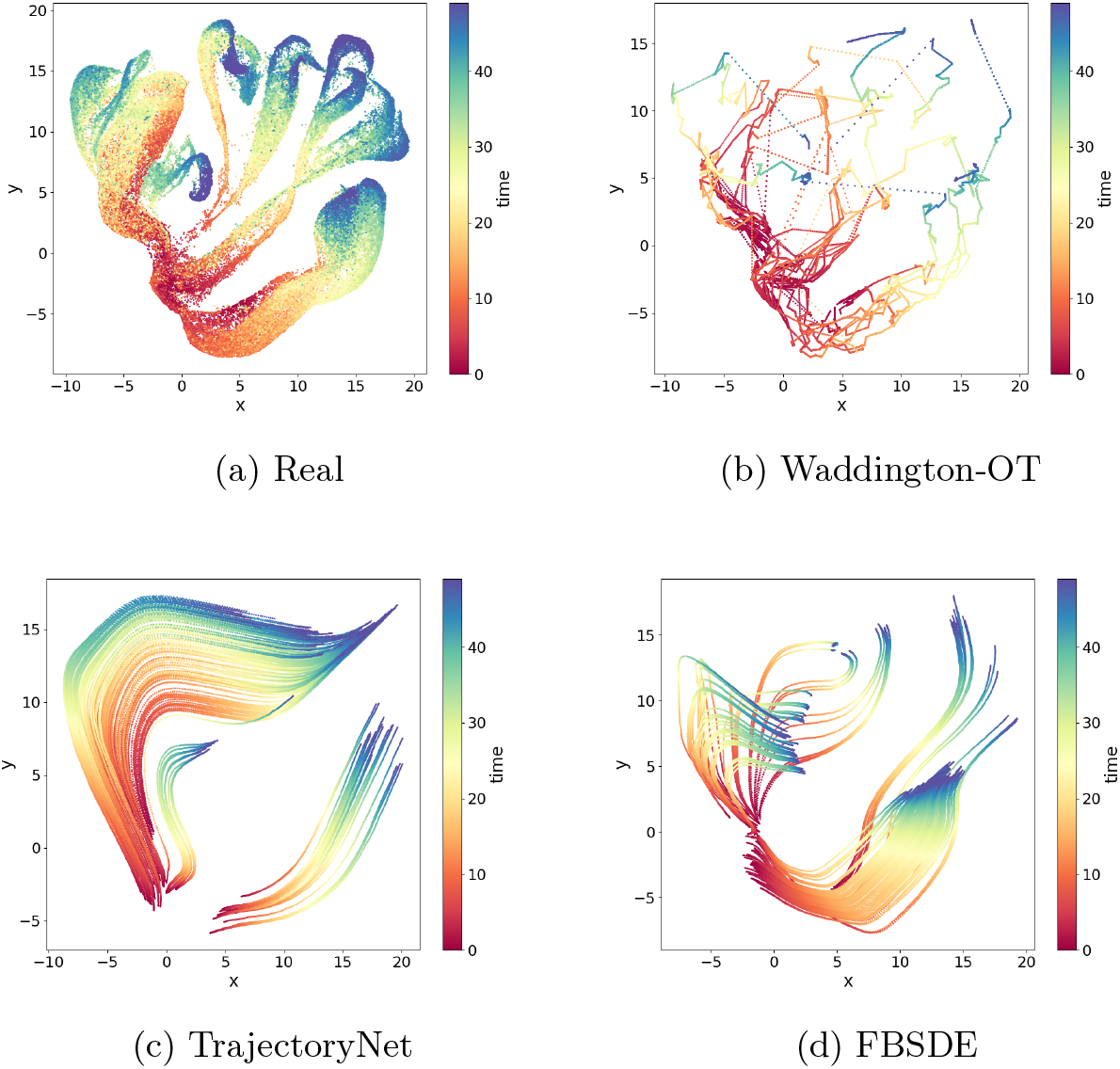
Performance on a *Arabidopsis thaliana* stem cells dataset. 6a) Gene expression of single cells are reduced to two dimensions using the UMAP embedding method. The data correspond to a total of 51 time points on a pseudotime scale from 0 to 50. 6b-6d) Trajectory inference by three different methods (Waddington-OT, TrajectoryNet and FBSDE). in each experiment, a total of 200 equally spaced time points are added between the starting and end time points. The same color code is used. As can be seen. Waddington-OT produces some unrealistic paths among different branches in 6a. TrajectoryNet fails to capture the geometric features of the true trajectories. FBSDE produces some trajectories that resemble the ground truth to some degree.

### 2.5 Cross-Validation Study

In this study, we performed a comprehensive benchmarking of the FBSDE method against two other prominent approaches, Waddington-OT (WOT) and TrajectoryNet. To evaluate their performances, we utilized cross-validation techniques on several well-established single-cell datasets.

For each validation set, we considered a total of *n* + 1 observed points labeled from *t*_0_ to *t*_*n*_. In the validation process, we randomly selected 20% of the time points as the test data, while the remaining time points were used for training. The model then interpolated the data from the selected 20% time points, as depicted in **Figure** 2, and compared the interpolated data with the ground truth (i.e., the test data). This process was repeated for a total of *M* = 50 times to ensure robustness, and the average performance metric across all repetitions was reported.

To assess the accuracy of interpolation, we chose the Wasserstein Distance *W*_1_ as the standard metric to measure the divergence between the interpolated data and the test data.

**Table** 1 summarizes the numerical performance for all three models on different datasets. For the human embryonic stem cells dataset, WOT generated distributions match the empirical distributions the best whereas FBSDE outperforms WOT and TrajectoryNet on the mouse embryonic fibroblasts dataset and the *Arabidopsis thaliana* stem cells dataset. While WOT simulations have higher precision (see definition of precision in **Figure** 2a), this is mainly due to the fact that WOT simulations are drawn from observations. On the other hand, when generative modelling is required for a larger number of time points such as in the mouse embryonic fibroblasts dataset and *Arabidopsis thaliana* stem cells dataset, the generative power of SDE models, i.e., TrajectoryNet and FBSDE, begin to outperform. Furthermore, our FBSDE also outperforms TrajectoryNet which is consistent from the visual comparison in **Figure** 4-6. In conclusion, the cross-validation study shows that, at least for the three datasets tested, our FBSDE model performs better than other contemporary methods in single cell trajectory inference.

**Table 1:** Performance Comparison.

|  | HES | MEF | ATS |
| --- | --- | --- | --- |
| Waddington-OT | 1.102(0.020) | 0.291(0.001) | 7.739(1.096) |
| TrajectoryNet | 1.202(0.002) | 0.287(0.001) | 6.182(0.004) |
| FBSDE | 1.189(0.006) | 0.177(0.011) | 5.808(0.006) |

The performance metric is the Wasserstein Distance $W_1$ with mean and standard deviation reported in brackets. HES: Human Embryonic Stem Cells Dataset. MEF: Mouse Embryonic Fibroblasts Dataset. ATS: *Arabidopsis thaliana* Stem Cells Dataset.

## 3 Discussion

In this paper, we present a new method called Forward-Backward Stochastic Differential Equations (FBSDE) to model the developmental trajectories of single cells and compare it to two other recent methods–Waddington-OT and TrajectoryNet. The FBSDE method has several advantages. First, the use of stochastic differential equations allows for the generative modelling of developmental trajectories on a continuous space-time manifold. Second, iterations between the forward SDE and the backward SDE converge to a solution that satisfies both the initial and terminal probability distribution constraints leading to higher accuracy in trajectory modelling. Third, the FBSDE method integrates concepts from mean field theory to mimic the effects of cell-to-cell interactions allowing for more realistic biological modelling.

We apply all three methods on several scRNA-seq data sets. In summary, the FBSDE method outperforms the other two methods significantly. In situations where accurate modelling is difficult to achieve (e.g., *Arabidopsis thaliana* stem cells dataset), the FBSDE method still retains a fairly high level of similarity to the ground truth.

Although FBSDE demonstrates great promise and success in generative modelling of single cell developmental trajectories, there exist a few limitations. Firstly, our model is applied to a two-dimensional dataset, which has been processed and dimensionally reduced from an initial dimensionality of up to 20,000 genes. While the model is promising in low dimensions, the curse of high dimensionality should not be overlooked in cases where a large number of genes cannot be reduced in a dataset. Recent works by Chen et al [27] and Liu et al [28] aim to address such a problem where each gene of each cell is modelled by a separate stochastic differential equation. Secondly, the interactions currently implemented in the FBSDE model do not cover a wide range of biological interactions. Despite their effectiveness, more complex interactions that model typical biological interactions such as ligand-signalling should be further investigated. Thirdly, it should be noted that the FBSDE model relies on the structure of the implemented neural network, and further exploration of different network architectures is warranted. Alternative forms of neural networks, such as the neural ODE proposed by Chen et al. [29] and recurrent neural networks, may have the potential to improve the accuracy of generative modelling and should be considered for future investigation.

Our FBSDE method introduces a novel aspect of modelling single cell developmental trajectories. There are a few improvements that can be made in future directions. First, instead of a single SDE, the FBSDE model can incorporate a mixture of stochastic differential equations analogous to Gaussian mixture models in probability density estimation. The introduction of the mixture model has the potential to increase model accuracy in situations such as the *Arabidopsis thaliana* stem cells dataset. Second, the FBSDE method can be generalized to systemic biology by incorporating spatial data into the genetic data. The introduction of spatial information allows for the modelling of cell-to-cell interactions that require physical proximity.

## 4 Methods

This section is structured as follows. In Section 4.1, we begin by providing a mathematical formulation of the developmental trajectory of a single cell and discussing the optimization problem typically employed in trajectory inference methods based on stochastic differential equations (SDEs). We also emphasize the importance of considering constraints within this optimization problem. Additionally, we present an overview of the TrajectoryNet framework [18], which serves as baseline for comparison.

Next, in Section 4.2.1, we address the approximation of the cost functional, which represents the optimal value of our optimization problem. The cost functional is defined as the expectation of a stochastic integral. To approximate this integral, we employ time discretization, also known as the Euler-Maruyama method. We further discuss the approximation of probability density functions (PDFs), which play a vital role in SDE-based trajectory inference. By implementing these numerical techniques, we illustrate how the cost functional can be effectively approximated given a finite sample.

In Section 4.2.2, we propose modifications to the TrajectoryNet framework to improve the approximation of the intermediate constraints. We provide detailed explanations of how these modifications can be implemented using a finite set of cells, leading to enhanced trajectory inference accuracy.

Finally, in Section 4.2.3, we introduce the Forward-Backward Stochastic Differential Equation (FBSDE) system and highlight its advantages in satisfying the terminal constraint. We implement a numerical algorithm proposed by Vargas et al. [21] for solving the FBSDE system.

### 4.1 Mathematical Details

Suppose all the cells live in the space of ℝ^*d*^ in the finite time frame [0, *T*] for some constant *T >* 0. The gene expression profile of a cell at time *t* is denoted by *X*_*t*_ *∈* ℝ^*d*^, which is a continuous random vector with probability density function *q*_*t*_ for *t ∈* [0, *T*].

#### Remark 1

In practical applications, gene expression data is often preprocessed and undergoes dimensional reduction to reduce its dimensionality. Typically, the reduced dimension *d* is much smaller, commonly set to 2 or 3. In our study, all the datasets were obtained in the form of two-dimensional data.

We are interested in the single-cell developmental trajectory *{X*_*t*_, *t ∈* [0, *T*} . Let *X*_0_ be an initial gene expression of a cell at time 0, then the trajectory of this cell at time *t* can be modeled by the stochastic differential equation as described by Zhang et al [12]:

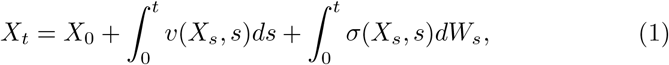

where *v* : ℝ^*d*^ *×* [0, *T*] is a drift function and *v*(*X*_*s*_, *s*) characterizes the shift change caused by *X*_s_ at time *s*. The integral 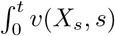 represents the cumulative mean shift induced by the trajectories of *X*_*s*_ up to time *t*, which is known as the drift term. The *σ* : ℝ^*d*^ *×* [0, *T*] is a volatility function and *{W*_*t*_*}*_0*≤t≤T*_ is a standard Brownian motion. At time *s, dW*_*s*_ refers to the change in the Brownian motion *W* . The term 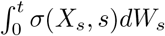 represents the random variation induced by the trajectories of *X*_*s*_ up to time *t*, which is known as the volatility term.

To infer the cell trajectory, we aim to learn the unknown drift function *v*(*X*_*t*_, *t*), also called action. In our analysis, we assume the volatility function is constant and set it equal to 1 as the default value.

**Equation** 1 characterizes the temporal dynamics of gene expression in an individual cell. In a population consisting of millions of cells, each cell undergoes simultaneous changes in its gene expression. Consequently, an action *v* governs the evolution of the population distribution over time. Remarkably, given an initial distribution *q*_0_, the sequence of distributions *ρ*(·, *t*)_0*<t≤T*_ can be mathematically represented as a function of the action *v* and volatility *σ* by leveraging the well-known Fokker-Plank Equation [30]. In this context, *ρ*(*·, t*) denotes the probability density function generated by the action *v* and satisfies *ρ*_*t*_ = *ρ*(, *t*) : ℝ^*d*^ × [0, *T*] → ℝ. We assume that *ρ* is a continuous function with continuous partial derivatives in the space ℝ^*d*^ × [0, *T*].

As cells undergo changes in their gene expression over time, there is often an associated energy cost. The optimal action 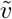 can be determined by seeking a function that minimizes this cost functional. This prompts the question of whether an optimal strategy exists for all cells in the population to collectively modify their gene expressions in order to minimize the cost functional. In general, most SDE-based inference methods consider minimizing the following cost functional:

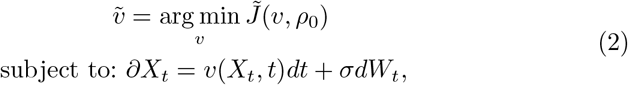

where

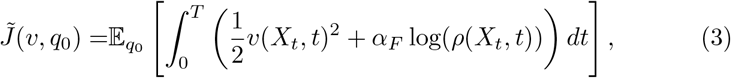

and *α*_*F*_ *>* 0 is a regularization constant tuned by cross-validation.

Since the initial gene expression *X*_0_ of a cell is a random variable with distribution *q*_0_, the cost functional is defined as the expected cost of a trajectory originating from all possible values of *X*_0_. The first term in the integral represents the non-interactive cost, where a cell’s trajectory is independent of the actions of other cells. The integral of this term is typically referred to as the kinetic energy of the trajectory, representing the amount of energy required to change a cell’s gene expression along this trajectory. In practice, the trajectory of one cell is likely to be influenced by other cells, therefore we need the second term in the integral, which is the running cost or interactive cost, representing how a cell is affected by other cells while performing action *v*. As a result, this term increases as the probability density at *X*_*t*_ increases. This concept is similar to the idea of crowdedness or entropy in terms of gene expression. For example, cells with similar gene expressions may compete for resources for growth, leading to trajectories with similar gene expressions having higher costs (i.e., higher *ρ*(*X*_*t*_, *t*)).

In trajectory inference settings [11, 31], one often observes *q*_*t*_ (or more precisely the empirical version of it) at a grid of time points 0 = *t*_0_, *t*_1_, *t*_2_, …, *t*_*K*_ = *T* . The trajectory inference problem aims to determine the optimal action 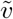 such that the population evolves from *q*_0_ to *q*_1_, *q*_2_, and ultimately to *q*_*T*_, while minimizing the cost functional specified in **Equation** 2.

Let *{ρ*_*t*_ : *t ∈* (0, *T*]*}*, denote a sequence of distributions generated from the initial distribution *q*_0_ through the application of the SDE model governed by the action *v*. In this context, the trajectory inference problem can be formulated as a constrained optimization problem,

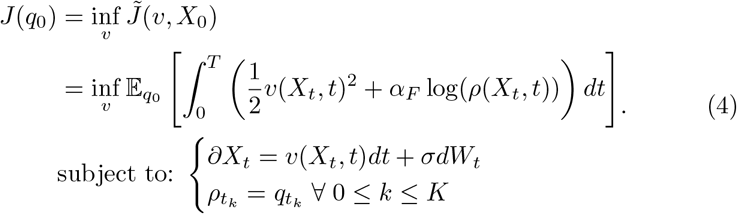

#### Remark 2

Consider the scenario where one aims to construct a road connecting two points *A* and *B*, while also passing through multiple intermediate checkpoints. In **Equation** 4, the SDE constraint *∂X*_*t*_ = *v*(*X*_*t*_, *t*)*dt* + *σdW*_*t*_ models the shape of the road. The initial, terminal, and intermediate constraints can be seen as analogous to the endpoints *A* and *B* and the intermediate checkpoints, respectively. Furthermore, the cost functional 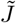 in the optimization problem corresponds to the cost associated with building the road. Thus, this optimization problem can be viewed as the task of designing a road that efficiently connects all the required checkpoints, while minimizing the overall cost 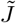.

However, previous studies [18, 21, 32] have argued that imposing equality constraints 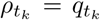 can be numerically challenging. These methods satisfy the initial constraint *ρ*_0_ = *q*_0_ by design. For all subsequent constraints at *t* = *t*_1_, *t*_2_, …, *T*, they approximate these equality constraints, denoted by 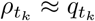. One example is TrajectoryNet which solves the following optimization problem:

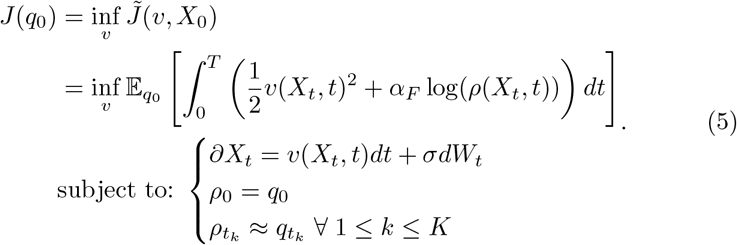

Ideally, the optimal solution to the relaxed problem in **Equation** 5 should be close to the optimal solution to the constrained problem in **Equation** 4. Nonetheless, the relaxed problem is prone to aggregation of errors. Moreover, it is important to note that the accuracy of approximating the constraint 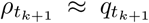 is dependent on the accuracy of approximating the previous constraint 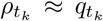. As a result, errors in approximation accumulate as *t*_*k*_ increases, potentially causing the terminal distribution *ρ*_*T*_ to deviate significantly from the true distribution *q*_*T*_ . This deviation can lead to erroneous information about the gene expressions of fully differentiated cells within the population. To mitigate this issue, we choose to enforce an exact terminal equality constraint *ρ*_*T*_ = *q*_*T*_ . By satisfying this constraint, we expect to improve the accuracy of approximations for the intermediate constraints. To fulfill the terminal constraint, we employ a Forward-Backward Stochastic Differential Equations (FBSDE) model proposed by Vargas et al [21]. In the following sections, we will describe how we integrate the TrajectoryNet framework and FBSDE to satisfy the terminal equality constraint and approximate the intermediate equality constraints. Specifically, our aim is to solve the following optimization problem:

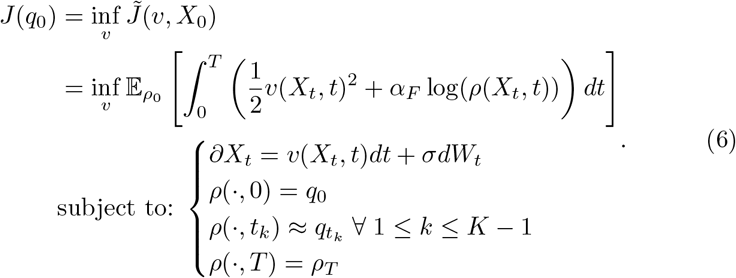

#### Remark 3

To better understand our model, imagine a string on a flat surface. The goal is to shape the string in a way that it connects two fixed points (initial and terminal constraints) while passing through specific points on the surface (intermediate constraints). TrajectoryNet holds one end of the string fixed while allowing the other end to roam freely, hoping to be close to the intermediate checkpoints and the terminal end. In contrast, our model fixes both ends of the string, increasing the likelihood of passing through the intermediate checkpoints and providing a better approximation of the intermediate constraints.

### 4.2 Implementation

In order to solve the optimization problem in **Equation** 6, we need to address the following numerical problems.

Firstly, we need to compute the cost functional [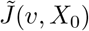, which involves calculating the expectation of a stochastic integral over continuous time with respect to the initial distribution *q*_0_. To approximate this integral, we adopt time discretization. By discretizing the SDE model on a dense set of time points between 0 and *T*, we can generate a finite number of simulated data to estimate the expectation. In particular, since the cost functional represents the expected cost of trajectories originating from initial points distributed according to *q*_0_, we sample *m* cells from the observations at *t* = 0 and employ a discrete-time approximation of the SDE model to simulate *m* trajectories.

Secondly, we need to estimate the true distribution *q*_*t*_ and the generated distribution *ρ*_*t*_ at all time points. This requires computing the probability density functions (PDFs) at the selected time points used in the stochastic integral approximation. To accomplish this, we employ kernel density estimation of the empirical distributions.

Thirdly, we need to approximate the intermediate constraints 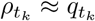 . In order to ensure that the simulated distributions closely match the empirical distributions at intermediate time points, we introduce a new penalty term along with the ones proposed by TrajectoryNet.

Lastly, it is crucial to satisfy the terminal equality constraint that ensures the agreement between the true distribution *ρ*_*T*_ and the simulated distribution *q*_*T*_ at the final time point *T* . To enforce this constraint, we incorporate the Forward-Backward Stochastic Differential Equations (FBSDE) model.

Additionally, in our notation, we assume the observations *X*_*t*_ are collected at *K* + 1 observed time points: 0 = *t*_0_ *< t*_1_ *< t*_2_ *<* … *< t*_*K−*1_ *< t*_*K*_ = *T* . At each time point *t*_*k*_, we have *n*_*k*_ observations, denoted by 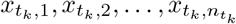.

#### 4.2.1 Estimation of Cost Functional

To estimate the cost functional, which represents the expectation of a stochastic integral, a discretization approach is employed. The integral of a function over continuous time is approximated by summing the function values multiplied by the interval lengths over numerous small intervals. Therefore, we need to generate a set of discrete time points for this purpose.

To be more specific, to account for the constraints imposed by the optimization problem in **Equation** 6 at the observed time points {*t*_*k*_}_0*≤k≤K*_, the set of discrete time points *U* is constructed by take a union of the observed time points *{t*_*k*_*}*_0*≤k≤K*_ and a denser set of equally spaced time points 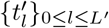 with 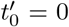 and 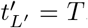. With a bit abuse of notation, we denote 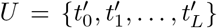, where 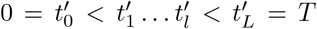. This incorporation of observed time points ensures that the discretization captures the necessary information at these specific instances.

##### Remark 4

For instance, if *T* = 1 and the observed time points are 0.33, 0.66, then the set *U* can be obtained by taking the union of *{*0, 0.05, 0.1, …, 0.95, 1*}* and *{*0.33, 0.66*}* and then ordering the numbers in the set in an increasing order.

Additionally, the expectation is approximated by averaging *m* simulated paths generated through the approximation of the stochastic differential equation (SDE) *∂X*_*t*_ = *v*(*X*_*t*_, *t*)*dt* + *σdW*_*t*_. The simulation process begins with the selection of an initial set of *m* cells with gene expressions 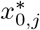 for 1 *≤ j ≤ m*, from the set of initial observations 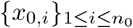 .

Based on the definition of *U* as the set of time points, the interval length 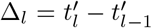 is assigned for 1 ≤ *l ≤ L*. The simulations are generated using the Euler-Maruyama scheme as follows [21]:

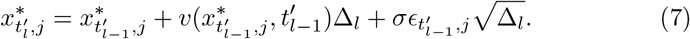

Here, 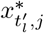 represents the gene expression of the *j*-th simulated cell at time 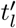 for 1 ≤ *j ≤ m* and 0 ≤ *l ≤ L*. The noise terms 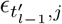 are independent standard normal random variables. **Equation** 7 approximates the constraint *∂X*_*t*_ = *v*(*X*_*t*_, *t*)*dt* + *σdW*_*t*_. It is important to note that the accuracy of the approximation in **Equation** 7 improves as the number of simulated time points *L* increases.

The outlined approximation scheme produces *m* simulated paths. In order to assess the stochastic integral, it becomes necessary to estimate the probability density function (PDF) *ρ*_*t*_ for each *t ∈ U* . Furthermore, it is essential to estimate *q*_*t*_ to ensure that the constraints *ρ*_*t*_ *≈ q*_*t*_ are satisfied at the observed time points *t*.

In particular, we can not obtain the true density functions 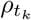 and 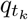, and we need to replace them by their estimates 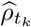 and 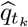 in solving the optimization problem in **Equation** 5. To estimate the true density function 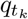 and 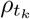, we use the Nadaraya-Watson density estimator. Based on the observations 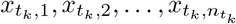, we estimate 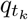 by

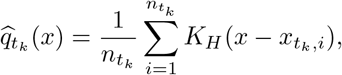

where *H* is a diagonal matrix with diagonal entries *h*. The bandwidth *h* is tuned by cross validation.

Similarly, based on the simulated points *{*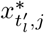: 0 *≤ l ≤ L*, 1 *≤ j ≤ m}*, we can estimate the distribution of the simulated points at time 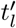 by

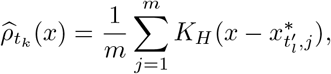

Again, the same kernel function *K*_*H*_ is used to compute the empirical distributions.

To summarize, we employ the discretized SDEs and perform kernel density estimations to simulate multiple trajectories and estimate the expected value of the stochastic integral. This allows us to approximate the cost functional. Specifically, for a given action *v*, the cost functional is computed as the average of the following terms over the simulated trajectories:

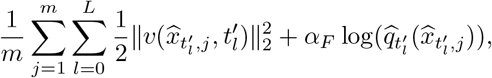

#### 4.2.2 Approximation of Intermediate Constraints

In order to approximate the intermediate constraints 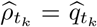, we introduce an additional penalty term along with the penalties proposed by TrajectoryNet [18]. The purpose of this new penalty term is to enhance the recall of the model, as demonstrated in **Figure** 2b.

We start by introducing some notations. Let

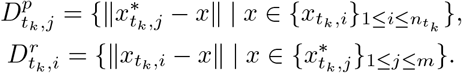

The set 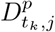 is the set of distances between the *j*th simulation at *t*_*k*_ and all observations at *t*_*k*_. The set 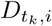 is the set of distances between the *i*th observation at *t*_*k*_ and all simulations at *t*_*k*_. Furthermore, we order both sets in ascending order such that 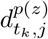 denotes the *z*th smallest element in 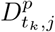 and 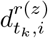 denotes the *z*th smallest element in 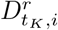 .

Subsequently, we implement the following penalty function *g*_*k*_ at observed time point *t*_*k*_,

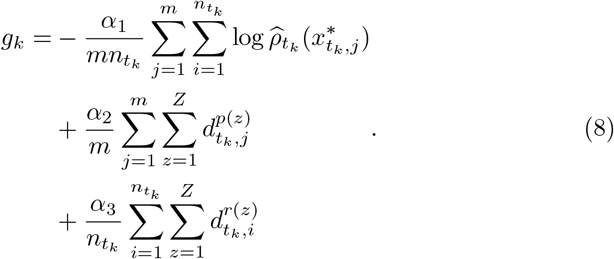

Here, *α*_1_, *α*_2_, and *α*_3_ are positive constants used for regularization.

The first two penalty terms in the penalty function are proposed by TrajectoryNet. The former term applies the maximum likelihood function to the simulations, computed from empirical distributions. It aims to encourage the simulations to closely match the empirical distribution of the observations at time *t*_*k*_. This term ensures that the simulations capture the statistical characteristics of the observed data.

The second term aims to improve the precision of the model by penalizing large distances between the simulated data points and observations. It pushes the simulations to be closer to the observed data points, promoting a better alignment between the simulated trajectories and the actual observations. By minimizing this penalty term, the model becomes more precise in capturing the observed trajectory patterns.

We introduce the third penalty term to enhance the recall of the model. It ensures that a larger number of observations are included in a close neighbourhood around the simulations. This term is necessary when the simulated data points are only close to a few observations, resulting in the first penalty term being close to zero, while the distribution of simulated data points, 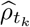, is not close to the empirical distribution 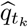 (see **Figure** 2a-2b).

With the combination of these three penalty terms, we aim to improve both precision and recall, leading to more accurate trajectory inference.

#### 4.2.3 Satisfying Terminal Constraint

This section presents a detailed implementation of Forward-Backward Stochastic Differential Equations (FBSDE) in order to achieve the terminal constraint 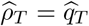 mentioned earlier.

In the continuous setting, an FBSDE model is a system of two stochastic differential equations where *X*_*t*_ is the forward process and *Y*_*s*_ is the backward process such that,

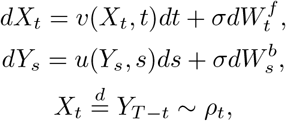

where 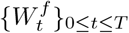 and 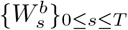 are two independent standard Brownian motions. The stochastic processes *{X*_*t*_*}*_0*≤t≤T*_ and *{Y*_*s*_*}*_0*≤s≤T*_ in ℝ^*d*^ with constant volatility *σ* are involved in the FBSDE model. The process *X*_*t* 0*≤t≤T*_ is equivalent to the trajectory expressed in **Equation** 1, describing the gene expression changes of a cell over time. The forward drift *v* specifies the current gene expression of a cell and current time to the direction in which the cell changes its gene expression at that time. The backward process *{Y*_*s*_*}*_0*≤s≤T*_ reverses the sequence *{X*_0_, *X*_1_, …, *X*_*T*_ *}*. In other words, if the distribution of gene expressions of fully differentiated cells is *ρ*_*T*_ at terminal time *T*, the initial distribution *ρ*_0_ at time 0 can be obtained by applying the backward process to the random variable *X*_*T*_ . Finally, the mathematical connection between the forward drift *v* and the backward drift *u* is stated in the following theorem.

##### Theorem 1

(Nelson’s Duality Theorem [33])

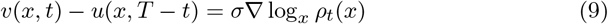

Theorem 1 establishes a relationship between the forward drift *v* for a cell with gene expression *x* at time *t* and the backward drift *u* for a cell with the same gene expression *x* at time *T − t*. Based on this relationship, Vargas et al. [21] proposed an algorithm to solve a simplified optimization problem described as follows:

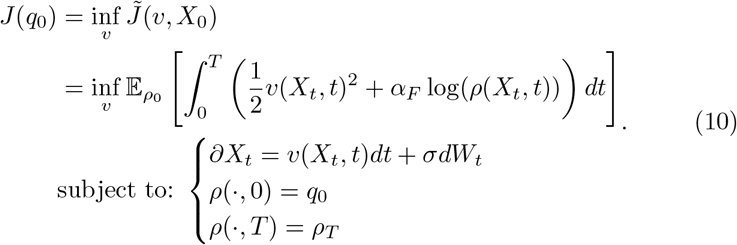

However, since this problem only considers the initial and terminal constraints, we need to incorporate penalty terms, as discussed in Section 4.2.2, to approximate the intermediate constraints presented in our main optimization problem described as follows:

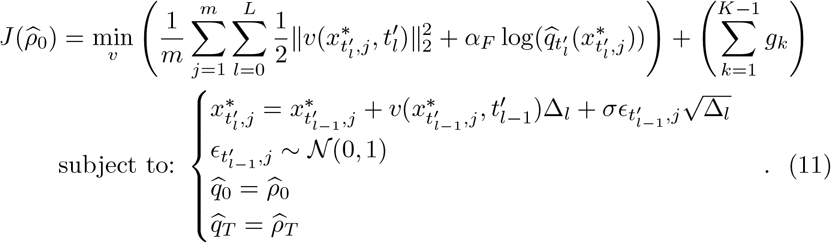

The algorithm proposed by Vargas et al. [21], as shown in **Figure** 1, consists of an iterative procedure involving two optimization problems: a forward optimization problem and a backward optimization problem. The forward optimization problem focuses on enforcing the constraint *ρ*_0_ = *q*_0_ and approximating the remaining constraints 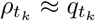 for all *k* = 1, 2, …, *K*. Conversely, the backward optimization problem focuses on enforcing the constraint *ρ*_*T*_ = *q*_*T*_ and approximating the constraints 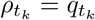 for *k* = 0, 1, 2, …, *K* 1. Both optimization problems incorporate a new penalty term. Specifically, the forward optimization problem penalizes the discrepancy between simulations generated by the forward drift and references generated by the solution to the backward optimization problem. Similarly, the backward optimization problem penalizes the discrepancy between simulations generated by the backward drift and references generated by the solution to the forward optimization problem.

The forward optimization problem is formulated as follows. Let *u*′ be the backward drift, and let 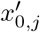 be the initial points sampled from observations *x*_0,*i*_. We define the *m* **reference** paths as:

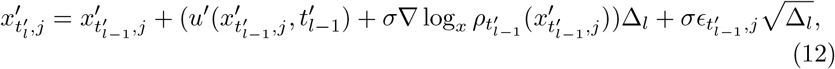

where 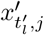 represents the *j*-th reference at time 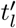.

The corresponding forward optimization problem becomes:

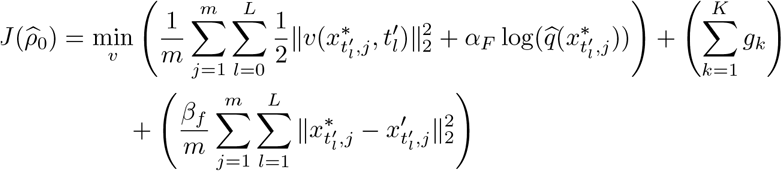

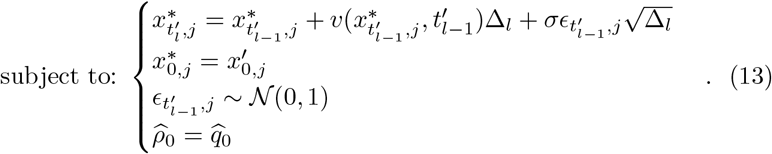

The corresponding forward optimization problem, denoted by **Equation** 13, aims to minimize the cost functional 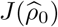 with respect to the action *v*.

Note that the objective function of the forward optimization problem includes an additional penalty term (with regularization constant *β*_*f*_ tuned by cross validation) to account for the divergence between the simulations 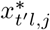 and the reference paths 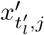. This term penalizes the differences between the trajectories generated by the optimal forward drift and the reference paths. Additionally, the terminal equality constraint 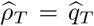 is converted into the penalty term *g*_*K*_ in the objective function.

In summary, the first three terms in the objective function are similar to the mean field problem described in **Equation** 11, with an additional term that quantifies the differences between the simulations generated by the optimal forward drift and the reference paths generated by the backward drift.

The backward optimization problem is defined as follows. Let *v* be the forward drift, and let 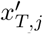 be the terminal points sampled from observations 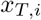. we define the *m* **reference** paths as:

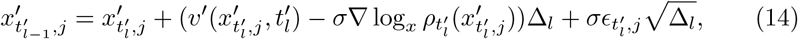

where 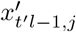 represents the *j*-th reference path at time 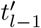.

The corresponding backward optimization problem becomes:

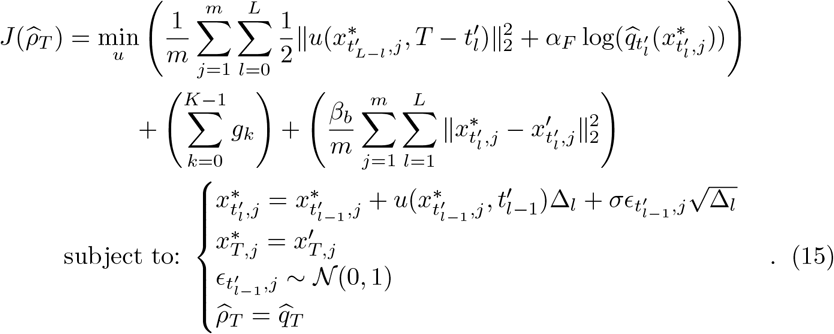

Similar to the forward optimization problem, the equality constraint 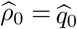 is converted into the penalty term *g*_0_. The objective function of the backward optimization problem includes the first three terms, which are the same as the mean field problem described in **Equation** 11. Additionally, there is an extra term that quantifies the differences between the simulations generated by the optimal backward drift and the references generated by the forward drift. This term captures the discrepancies between the backward simulations and the forward references.

To summarize, the proposed algorithm by Vargas et al. [21] iterates between the forward and backward optimization problems, where the optimal forward drift generates references for the computation of the optimal backward drift, and vice versa. In each iteration, penalty terms are used to penalize the differences between simulations generated by the optimal drift in one direction and references generated by the drift in the opposite direction. The algorithm has been shown to converge [21, 22, 24, 32], with the optimal forward drift satisfying the terminal constraint 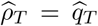 and the optimal backward drift satisfying the initial constraint 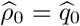. This implies that the optimal forward drift obtained from this iterative process is indeed the solution to the original optimization problem in **Equation** 11, satisfying both the initial and terminal constraints.

To prepare for the computational algorithm, we will parameterize the drift functions *v* and *u* using neural networks that will be denoted as *ϕ* and *ψ*, respectively. Specifically, we will initialize the parameters in these neural networks so that they can be trained to approximate the desired drift functions. Both neural networks will be fully connected, with default setting of three hidden layers each containing 128 units, and they will use ReLU activation functions throughout.

##### Algorithm 1 Solving FBSDE Model in **Equation** 11

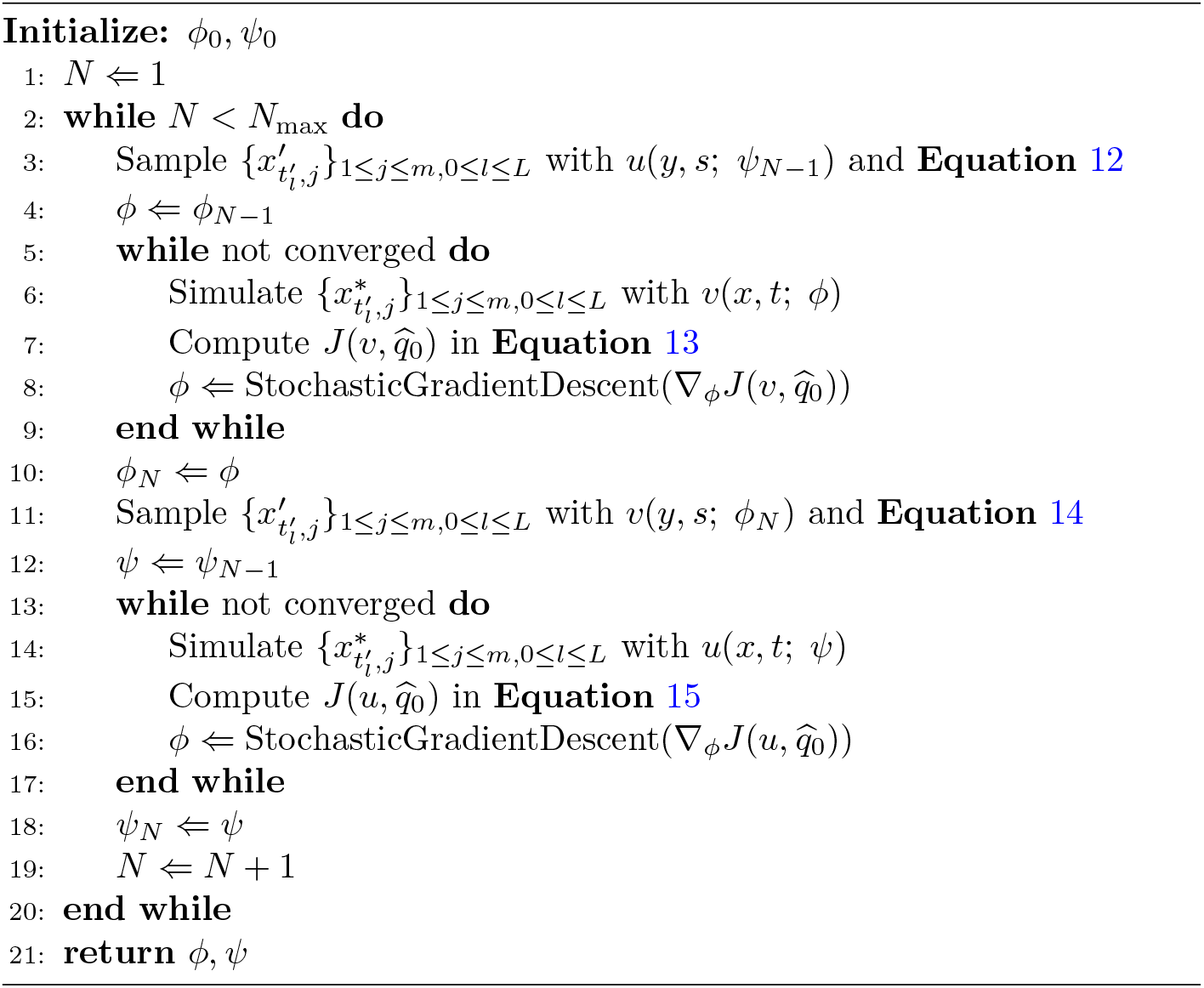

## Declarations

### Funding

DK and ZZ acknowledge financial support from a Catalyst Grant from Data Science Institute and Medicine by Design, University of Toronto. KZ and JZ is partially supported by CANSSI (Canadian Statistical Sciences Institute).

### Conflict of interest/Competing interests

None.

### Ethics approval

Not applicable.

### Consent to participate

Not applicable.

### Consent for publication

All the authors have read the manuscript and approved the submission at the current form.

### Data availabilty

– The Human Embryonic Stem Cells dataset provided by Moon et al is available from the Mendeley Data repository at https://doi.org/10.17632/v6n743h5ng.1.
– The Mouse Embryonic Fibroblasts dataset provided by Schiebinger et al is available from NCBI Gene Expression Omnibus at https://www.ncbi.nlm.nih.gov/geo/query/acc.cgi?acc=GSE122662.
– The*Arabidopsis thaliana* Stem Cells dataset provided by Shahan et al is available from NCBI Gene Expression Omnibus at https://www.ncbi.nlm.nih.gov/geo/query/acc.cgi?acc=GSE152766.

### Code availability

All code used in this study are available from the GitHub respository at https://github.com/Diebrate/population_model.

### Authors’ contributions

– Conceptualization: DK, ZZ
– Data curation: KZ
– Methodology and investigation: KZ, JZ, KZ, ZZ
– Writing – original draft: KZ, DK, ZZ

## Notes

https://doi.org/10.17632/v6n743h5ng.1

https://www.ncbi.nlm.nih.gov/geo/query/acc.cgi?acc=GSE122662

https://www.ncbi.nlm.nih.gov/geo/query/acc.cgi?acc=GSE152766

### Competing Interest Statement

The authors have declared no competing interest.

